# Machine Learning to Predict Gut Microbiomes of Agricultural Pests

**DOI:** 10.1101/2024.08.12.607564

**Authors:** Md Jobayer, Alexander Taylor, Md Rakibul Hasan, Khandaker Asif Ahmed, Md Zakir Hossain

## Abstract

**Context:** While current efforts to control agricultural insect pests largely focus on the widespread use of insecticides, predicting microbiome composition can provide important data for creating more efficient and long-lasting pest control methods by analysing the pest’s food-digesting capacity and resistance to bacteria or viruses.

**Aims:** Instead of using computationally expensive techniques, we aim to investigate the dynamics of these microbiome compositions using metagenomic samples taken from fruit flies.

**Methods:** In this paper, we propose the three machine learning-based biological models. Firstly, we propose the *intrafamilial successor prediction*, which predicts the relative abundance of each bacterial family using the past four generations. Next, we propose our *interfamilial quantitative prediction*, where the model predicts the amount of a given bacterial family in each sample using the amount of all other bacteria present in the sample. Lastly. we propose our *interfamilial qualitative prediction*, which predicts the relative abundance of each bacterial family within a sample using binary information of all bacterial families.

**Key Results:** All three models were tested against Least Angle Regression, Random Forest, Elastic-Net, and Lasso. The third approach exhibits promising results by applying a Random Forest with the lowest mean Coefficient of Variance of 1.25.

**Conclusion:** The overall results of this study highlight how complex these dynamic systems are and demonstrate that more computationally efficient methods can characterise them quickly.

## 1 Introduction

As the global population continues to grow and food security issues increase in importance, future efforts to improve productivity and, ultimately, the yield of crops must also intensify Savary et al. (2019). A critical element of this effort is the management of and response to biophysical threats that can potentially cause devastating crop losses Sundström et al. (2014). These threats can be grouped into three distinct categories: environmental degradation, climate change, and pests and diseases of animals and plants Sundström et al. (2014). As threats within and between groups are often strongly correlated, it is difficult to comprehensively formulate potential management strategies in isolation Reddy et al. (2019).

Among crops, one of the most significant threats is insect pests, which reduce the amount of helpful produce and account for losses ranging from 10% to 40% worldwide Savary et al. (2012). Typically owed to their large population sizes, short generation times and high fecundity, these pests can rapidly change their abundance and distribution in response to a range of environmental pressures Pareek et al. (2017). Furthermore, as the effects of climate change begin to intensify and temperatures rise in tropical and subtropical regions where a range of agriculturally important crops are grown, the geographic range in which these insect pests can span is expected to expand significantly Sultana et al. (2020).

Current efforts to reduce the impact that these insect pests have on crops are typically focused on the widespread use of insecticides Pavlidi et al. (2018) due to their relatively low cost, their ease of application to crops, and the diversity of insect pests they can control Aktar et al. (2009). Despite these advantages, the unchecked application of insecticides presents a wide range of consequential issues, such as the nonspecific killing of beneficial insects Hill et al. (2017), the emergence of resistance within target pests Vontas et al. (2011), heavy environmental pollution Zhang et al. (2011) and, in some cases, negative impacts on human health Igbedioh (1991). As a result, many emerging management strategies have started to focus on using more targeted, cheaper insecticide alternatives that can sustain higher efficacy over much longer time frames Furlan et al. (2021).

Some common alternatives include the integration of transgenic genes into plants Schnepf et al. (1998), the introduction of beneficial predators and parasites Caltagirone and Doutt (1989), the attraction and trapping of insect pests using pheromones Kirsch (1988), and the application of natural, horticultural oils Nile et al. (2019). While these alternatives have been proven to be efficacious in varying degrees, various financial, societal and environmental drawbacks limit their use to specific applications Aidley (1976). One relatively novel and particularly interesting alternate strategy is the development of a highly specific bio-agent that can modulate the pest behaviour and has the potential to be utilised as biopesticides.

The gut microbiome is defined as the collection of microbial species that colonise the gut of a host organism Little and Light (2022). This microbiome plays a significant role in many biological functions of the host organism, including salvaging energy from otherwise undigestible nutrients Hooper et al. (2002), assisting the immune system Hooper and Gordon (2001), and even regulating host fat storage Palmer et al. (2007). Microbiome composition can influence insect control by influencing the insect’s capacity to digest food, resist bacteria and viruses, and interact with the environment Ami et al. (2010); Kyritsis et al. (2017). Researchers can learn more about how they might be able to modify the microbiome to control the behaviour or population of the insect by predicting the microbiome composition of a particular insect species Steinigeweg et al. (2022). For example, suppose a specific insect pest has a microbiome necessary for its own survival. In that case, researchers might disrupt the microbiome using targeted microbial therapies (such as probiotics or antibiotics) to lower the pest population by generating stronger host sterile males, competing better against wild flies. Similarly, if the microbiome impacts an insect’s food habits, researchers may be able to utilise this knowledge to create new insecticides that target specific microorganisms in the insect’s stomach Steinigeweg et al. (2022). Overall, predicting microbiome composition can give helpful information for building more effective and sustainable pest control strategies.

While in most cases, maintaining a healthy gut microbiome is the goal, the deleterious effects that come from imbalances present an opportunity for targeted control of certain species to alter the biochemistry and behaviour of the host Broderick et al. (2014). While metagenomic tools can accurately identify the exact composition of a small number of gut microbiomes, the computational complexity required to process the large datasets that describe vast numbers of samples is still too high Tang et al. (2013). One potential method of characterising and predicting the composition of a large number of gut microbiomes in a more computationally affordable yet coarse-grained way involves the use of machine learning (ML) techniques to make several simplifying assumptions Herńandez Medina et al. (2022). The premise that underpins the application of ML methods in this field is their ability to identify patterns in large sets of data without being explicitly programmed to do so Herńandez Medina et al. (2022). From these patterns, simplified assumptions can be made to construct models that can estimate the microbiome of a sample based on several input parameters.

The gut microbiome can be predicted using various methods, including abundance-dependent methods such as Analysis of Composition of Microbiomes (ANCOM) Mandal et al. (2015) and distance-based methods like Similarity Percentage Analysis (SIMPER) Schulz et al. (2021). However, these methods have their limitations. For instance, ANCOM performs poorly when the abundance of Operational Taxonomic Units (OTUs) exceeds 25% or is very low Mandal et al. (2015); Khomich et al. (2021). On the other hand, SIMPER only allows pairwise comparison, which can lead to misleading statistical output due to variance in the number of OTUs Mandal et al. (2015). Additionally, mathematical models Inamine et al. (2018) are highly specific to the host type of the microbiome Kumar et al. (2019). For example, if a model is built for a human host-based microbiome, we need to make a hypothesis and test it in-vitro and in species other than humans to make it a generic model. High dimensionality also poses a problem by increasing computational complexity. In this paper, we implement models that overcome the limitations of context-specific models and high dimensionality by focusing on the most valuable features of genomic data.

There are around 3000 fruit flies worldwide and Queensland fruitfly (Qfly) is one of the major horticultural pests in Australia. While there are many studies on these fruit flies gut microbiome and microbiome-regulated behaviours, no machine learning (ML) model has been developed so far to smoothen the microbiome study and assist in agricultural pest management. This article has several contributions to fill these gaps. At first, we use metagenomic samples taken from fruit flies (tephritidae) to analyse their microbiome composition. Using this knowledge, we observe how these microbiome compositions change with the developmental stage of the fly, as well as how they change across flies successively bred within a laboratory. Secondly, we analyse whether the complete microbiome composition of fruit flies or elements within this microbiome can be predicted by different ML models, using data obtained from experiments as predictors. Thirdly, we evaluate which ML models are best suited to predict systems such as gut microbiomes. The results of this research are intended to be used as tools to identify vital taxological families, genera and species of the microbiome that could be selected as targets for future insect pest control methods and to provide preliminary indications of the microbiome composition of a given sample.

Studies investigating the microbes present in gut samples usually originate from flies caught in the wild or reared in a laboratory. In either case, metagenomic samples are typically collected from whole flies or dissected guts of surface-sterilised flies. After DNA extraction and amplification of bacterial-specific sequences, the resulting samples were sequenced in a high-throughput sequencing platform. Within the literature, it is apparent that the most common bacterial families present in the gut microbiome of wild fruit flies are *Enterobacteriales* and *Lactobacilliales*. These bacterial families’ ubiquity is mainly due to the probiotic effects they confer on the host flies Kyritsis et al. (2019); Matos and Leulier (2014). An interesting study from Bakula (1969) demonstrated a strong link between the microbiome and developing larvae, indicating that a reduction in the amount of *Enterobacteriales* present can significantly increase the larval period. These observations were followed up by Ridley et al. (2012) where they noted that a reduction in the amount of *Acetobacter* also increased the larval period, but also significantly reduced the metabolic rate of flies, altered their carbohydrate allocation and increased glucose levels within the blood.

On the other hand, ML models have become increasingly popular for predicting gut microbiome-based outcomes. It is evidenced by a significant increase in publications on the gut microbiome’s role in colorectal cancer, which has risen 200 times from 2000 to 2021 Yu et al. (2023). Regarding ML-based microbiome prediction, Yang et al. (2021) conducted a study to investigate how the microbial community of black soldier fly changes over time due to host starvation. They observed a significant decrease in community diversity during the experiment. Using the Random Forest machine learning algorithm, they identified the most influential features that predict changes in the community. Notably, features such as ‘Cell growth and death,’ ‘Transport and Catabolism,’ and ‘Cancers’ were among the top predictors.

Also, an experiment was conducted by Turner et al. (2022) to determine the correlation between wild fish and their gut microbiota. Two species of teleost wild fish were collected from multiple lakes and islands, and the locations were categorised into two groups based on anthropogenic aquatic impacts: compromised aquatic environments (CAE) and non-compromised aquatic environments (non-CAE). Using ML models, the researchers attempted to classify new fish of those species into either CAE or non-CAE categories.

Apart from that, Pietrucci et al. (2020) describes the first ML data analysis of Parkinson’s disease microbiota dysbiosis using Random Forest, Neural Network, and Support Vector Machine models. However, the discrepancies in their outcome were attributed to a lack of standardised experimental and bioinformatic protocols, and the authors recommended creating a unique standard to ensure reliable comparisons in future studies. Other ML-based microbiome predictions include an experiment conducted on 2,320 individuals from Prince of Wales Hospital in Hong Kong. The author utilised their fecal microbiome sequence data to predict nine well-characterised phenotypes Su et al. (2022). Another study by Oh and Zhang (2020) created a learning framework called DeepMicro that utilises Autoencoders and other ML algorithms to classify diseases such as inflammatory bowel disease, type-2 diabetes and colorectal cancer. Their proposed framework outperforms the current state-of-the-art approaches conducted on humans while significantly reducing the input data’s dimensions.

## 2 Methodology

### 2.1 Data Description

We used data from Majumder et al. (2022) to construct the ML models. The raw data is presented in a tabular form that contains the total measured value of specific OTUs, which can then be mapped to particular bacterial phyla, classes, orders, families and genera. Within the raw data, there were 452 OTUs and 115 samples. After concatenating the OTUs to their corresponding bacteria, there were 6 unique phyla, 13 unique classes, 13 unique orders, 21 unique families and 47 unique genera.

Each sample represents a fly mapped to a developmental stage and generation using a provided mapping file. The stages include larvae, pupae, and adult flies, with adults further divided by male or female sexes. Samples are categorised into one of five generations, with most sets consisting of six samples. Sample size details can be found in the supplementary material (Table 4).

To simplify ML models, OTUs were grouped by taxonomic families instead of genera, considering that variation is prominent at the family level Majumder et al. (2022). The raw data lacked units for bacterial measurements, so normalisation was performed. The relative abundance of each OTU was calculated as a fraction of the bacterial family’s amount relative to the total bacteria within each sample. This normalisation created composition vectors for each sample, summing up to 100%.

To assess the quality of the processed data, the variance in relative abundances of bacterial families was measured for each developmental stage. Correlation matrices were generated to understand the pair-wise correlation between bacterial families within each stage and across all samples. An average matrix was constructed from the previous correlation matrices to identify trends in these correlations.

### 2.2 Model Development

#### 2.2.1 Model 1 – Intrafamilial Successor Prediction

This model, as shown in Fig. 1, aims to predict the relative abundance of each bacterial family in Generation 5 using the past four generations as predictors. The ‘n’ refers to the number of samples in each generation. The first four generations of each developmental stage were grouped to form the predictor set, while the generation 5 samples formed the outcome set. As each developmental stage has a different number of samples in Generations 1 to 4, each was reduced to the smallest size, resulting in sizes of n = 3 for Generation 1, n = 5 for Generation 2, n = 6 for Generation 3, n = 6 for Generation 4 and n = 6 for Generation 5. Consequently, this resulted in training data with a predictor set of size n = 20 and an outcome set size of n = 6 for each bacterial family in each developmental stage.

**Fig. 1.**
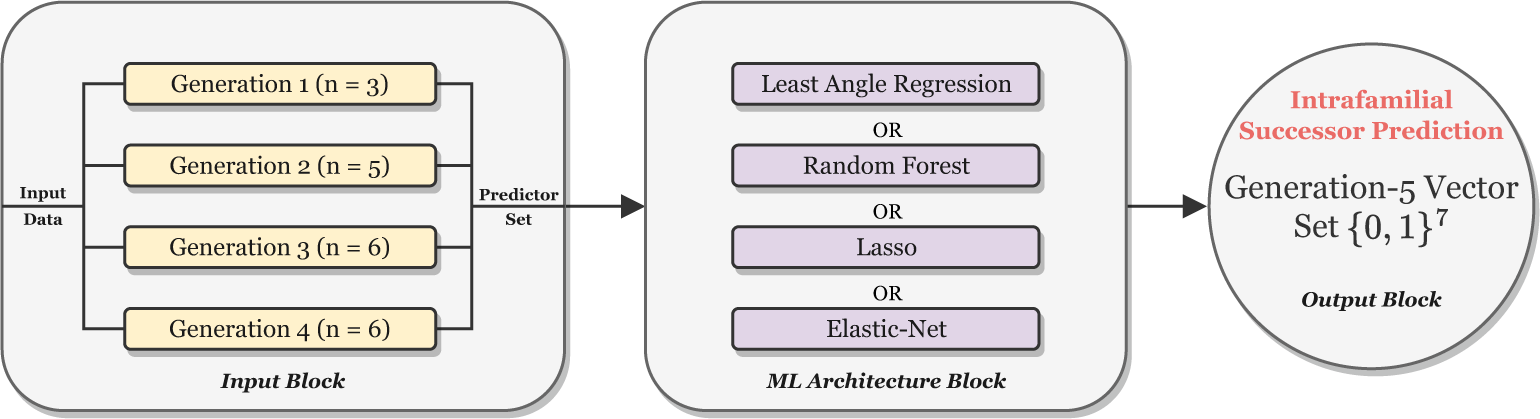
Intrafamilial Successor Prediction model architectural overview. Here, ‘n’ refers to the number of samples in each generation.

Four different ML methods were employed to construct models: Least Angle Regression, Random Forest, Lasso and Elastic-Net. The model was then tested on the adult Male developmental stage samples to measure the performance. The root mean square error (RMSE) was selected as the evaluation metric for the model due to its low computational cost and simplicity of implementation Wang and Bovik (2009).

Leave-one-out cross-validation was employed to obtain a more robust RMSE estimation and detect and prevent data over-fitting. From this cross-validation process, the standard error of measurement for the test scores was calculated so that it could be used to compare the performance between the different ML methods.

#### 2.2.2 Model 2 – Interfamilial Quantitative Prediction

This model aims to predict the amount of a selected bacterial family in each sample using the amount of all other bacteria present in the sample as predictors (Fig. 2). Because this model uses data containing the amount of other bacterial families as predictors, the relative abundances cannot be used, as this would reduce the model to a simple subtraction of all relative abundances from 100%. Consequently, the raw measurements provided in the data were used to construct and test the ML models.

**Fig. 2.**
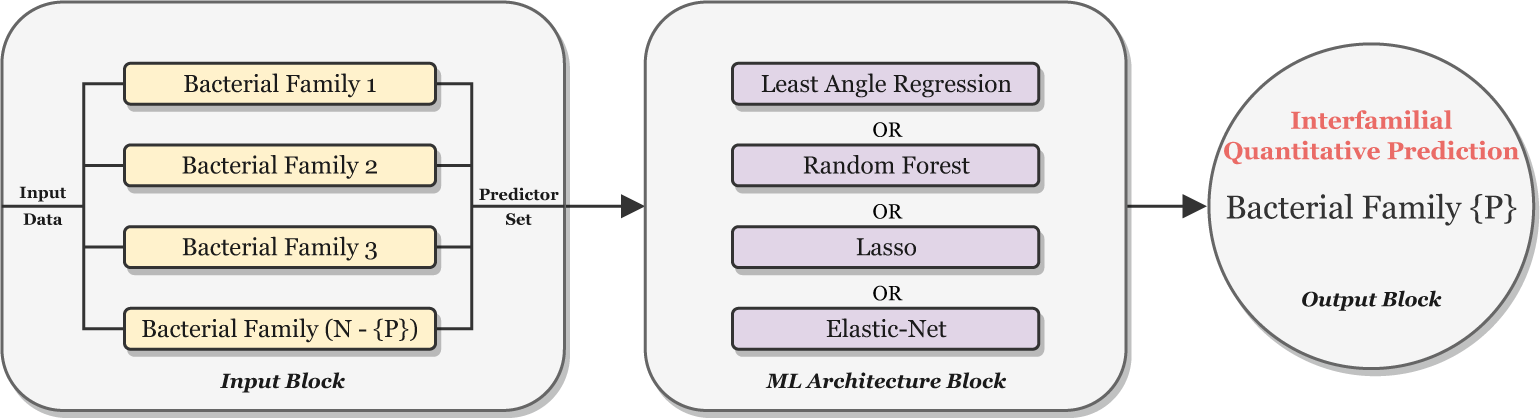
Interfamilial Quantitative Prediction model architectural overview. We use the variable ‘N’ to denote the total number of bacterial families to represent a generic figure. Additionally, we assign the label ‘P’ to the particular bacterial family that the model will predict, and we represent it as an element of a set enclosed in curly braces.

The dataset was iterated over, whereby one bacterial family was selected as the outcome set for each iteration, while all other bacterial families formed the predictor set. Furthermore, the same four ML methods employed for constructing Model 1 were used again. The model was then tested to measure the performance, with the Coefficient of Variance (CV) selected as the test score with leave-one-out cross-validation. It was done, as CV is a standardised form of RMSE, being normalised by the mean of the actual values, which allows greater comparison between different bacterial families Brown (1998). The mean CV and standard error were calculated from this set of CV values obtained through the leave-one-out cross-validation.

Families of several orders, such as the Lactobacillales Yoon et al. (2021), Enterobacteriales Sivakala et al. (2021), and Bacillales Glasl et al. (2019), are abundant in the gut microbiome of fruit flies and are linked to certain roles in the microbial communities. Such behaviour facilitates the formation of co-occurrence patterns among the families, therefore projecting the interfamilial quantitative predictive model’s capability.

#### 2.2.3 Model 3 – Interfamilial Qualitative Prediction

The construction of this model was adapted from Michel-Mata et al. (2022) and was developed in three phases. The first phase aimed to predict the relative abundance of each bacterial family using information on the presence or absence of all bacterial families within a sample as predictors. To create the predictor set for this model, each corresponding *composition vector* for the samples was reduced to a family vector, *P ∈* {0, 1}*^N^*. These vectors contain binary elements, where the *ith* bacterial family, *P_i_*, is assigned to 1 if present and 0 if absent. The original *composition vectors* were used as the outcome set. The model was then tested to measure the performance, with RMSE selected as the test score and leave-one-out cross-validation employed. From this set of RMSE values obtained through the leave-one-out cross-validation, the mean RMSE and standard error were calculated.

The second phase individually introduced two new predictor sets into the model. The first set contained *generation vectors*, *G ∈* {0, 1}^5^, describing which of the five generations the sample belonged to. The second set contained *developmental stage* vectors, *D ∈* {0, 1}^4^, indicating whether the sample was from a larvae, pupae, adult female or adult male fly. These vectors were individually appended to the family vectors, and the data was split into a training and testing set in the ratio of 70:30. The model was then tested to measure the performance, with RMSE selected and leave-one-out cross-validation employed. The third phase appended all three predictor vectors with the same method of training and testing applied. Finally, we obtain our desired set of bacterial family vectors by following the cNODE deep learning architecture Michel-Mata et al. (2022). A pictorial representation has been shown in Fig. 3.

**Fig. 3.**
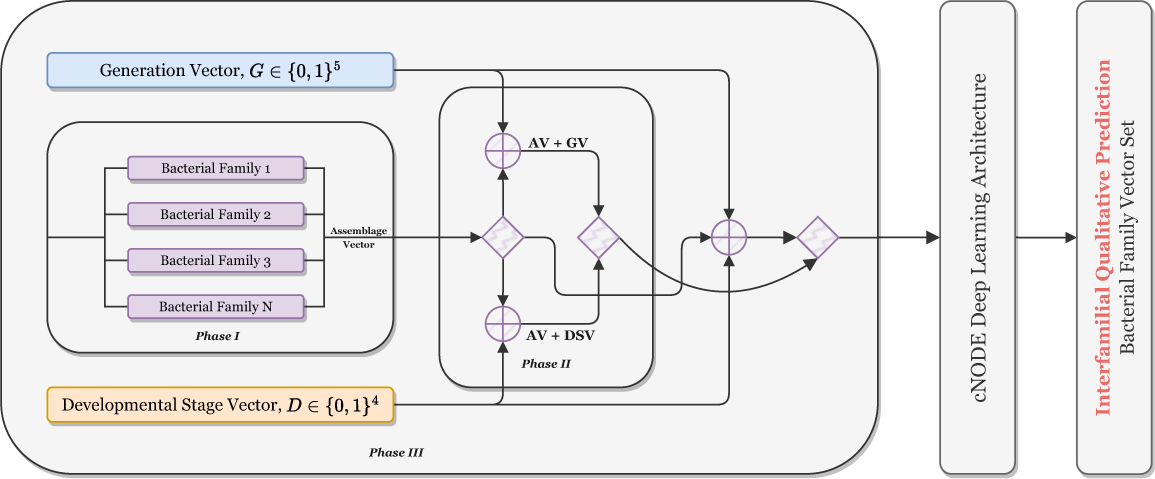
Interfamilial Qualitative Prediction model architectural overview. This model is comprised of three distinct phases. Phase I’s input is comprised of bacterial families of four generations. In phase II, the input is comprised of the input of phase I cascaded with generation vector G and developmental stage vector D separately. Unlike phase II, the input data in phase III comprises the input values of phase I and both generation vector G and developmental stage vector D simultaneously. The output or target column remains the same in all phases: the Generation 5 vector. Here, cNODE refers to the ‘compositional neural ordinary differential equation’ deep learning architecture proposed in Michel-Mata et al. (2022).

This qualitative predictive ML model has an advantage over the traditional approach in using quantitative microbiome profiles like species-level relative abundances and strain-specific markers to generate accurate predictionsPasolli et al. (2016). Moreover, this kind of ML model can forecast complex relationships between the microbiota and their host, providing insights into the interactions. Gould et al. (2018).

## 3 Results

### 3.1 Data Variance and Correlation

To inform the development of the ML models, the underlying data was characterised to understand the levels of variance amongst the number of bacteria present and identify any high-level correlation that may exist between the relative abundance of different bacterial families. The larvae samples exhibited the highest diversity in the measured number of families, with 16 different families present in this developmental stage, consistent with the findings in Majumder et al. (2022). As a result, the variance in the relative abundances of families was observed to be the largest in larvae samples. Different types of variance in the larvae samples are highlighted in Fig. 4a, describing the relative abundance of a bacterial family.

**Fig. 4.**
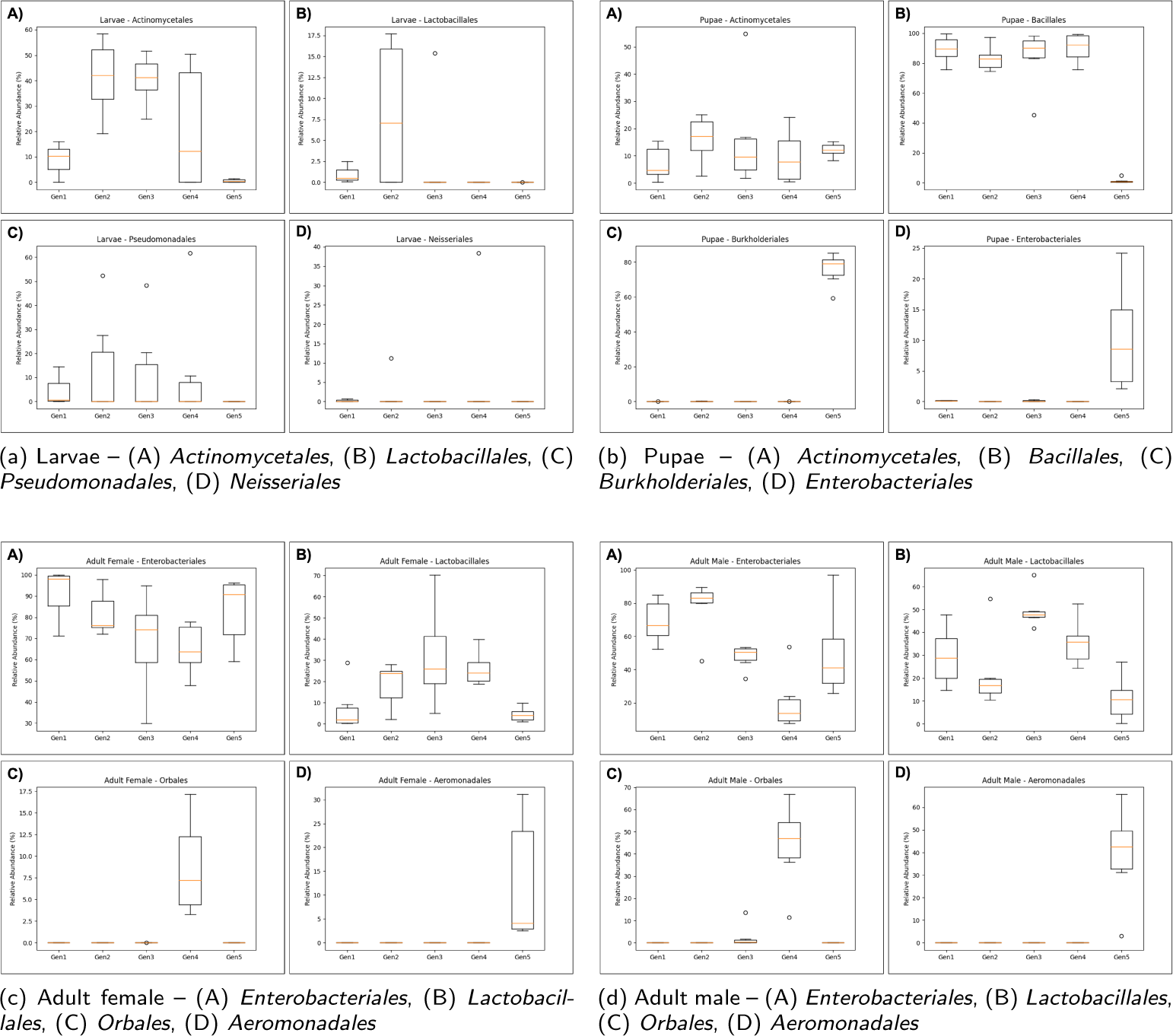
Relative abundances of some selected bacterial families ranging from Generation 1 to 5.

Fig. 4a (A) provides an example of the large variance in the abundance of a family observed across different generations, with the amount of *Actinomycetales* sharply increasing between Generation 1 and 2, remaining steady across Generation 2 and 3, and then sharply decreasing between Generations 3 to 5. Conversely, Fig. 4a (B) demonstrates how a bacterial family, such as *Lactobacillales*, can be almost entirely absent from every generation but present in a relatively significant proportion in a single generation. For some families, such as *Pseudomonadales*, the mean abundance across all generations is almost zero; however, a handful of samples skew the spread of measured abundance towards large values, as highlighted in Fig. 4a (C). Finally, several bacterial families are almost entirely absent from every generation in the larvae developmental stage. Yet, one or two samples record a large amount of the bacteria, as observed with *Neisseriales* in Fig. 4a (D).

The lowest diversity among all the families was observed in pupae samples, with 12 different families present in this developmental stage. Consequently, the lowest variance was observed in pupae samples, as evidenced in Fig. 4b, describing the relative abundance of four bacterial families. The majority of the bacteria present in pupae samples from Generations 1 to 4 are *Actinomycetales* and *Bacillales*, as shown in Fig. 4b (A) and 4b (B). In Generation 5, the amount of *Bacillales* drops significantly, being largely replaced by *Burkholderiales* and *Enterobacteriales*, as shown in Fig. 4b (C) and 4b (D). The other eight families observed in the pupae developmental stage are present in almost insignificant amounts (*<*3%) for every sample of each generation. Similarly, the changes in the adult female and male developmental stages are shown in Fig. 4c and Fig. 4d, respectively.

Fig. 5 shows that most bacterial families exhibit significant variations and low correlation among them. Handling such variations requires careful feature engineering and preprocessing to manage differences in scales, distributions, and missing values. A correlation heatmap was generated to identify high-level relationships between the families. Notably, *Bacillales*, *Lactobacillales*, *Enterobacteriales*, and *Actinomycetales* show significant relationships (Fig. 5). However, clusters of high correlation, particularly among *Erysipelotrichales*, *Bacteroidales*, and *Clostridiales*, are often due to their low relative abundance. The strongest relationship is observed between *Bacillales* and *Enterobacteriales*, with a correlation coefficient of −0.66, indicating antagonistic relative abundances. Similarly, *Actinomycetales* and *Enterobacteriales*, as well as *Bacillales* and *Lactobacillales*, exhibit negative correlations of −0.55 and −0.42, respectively.

**Fig. 5.**
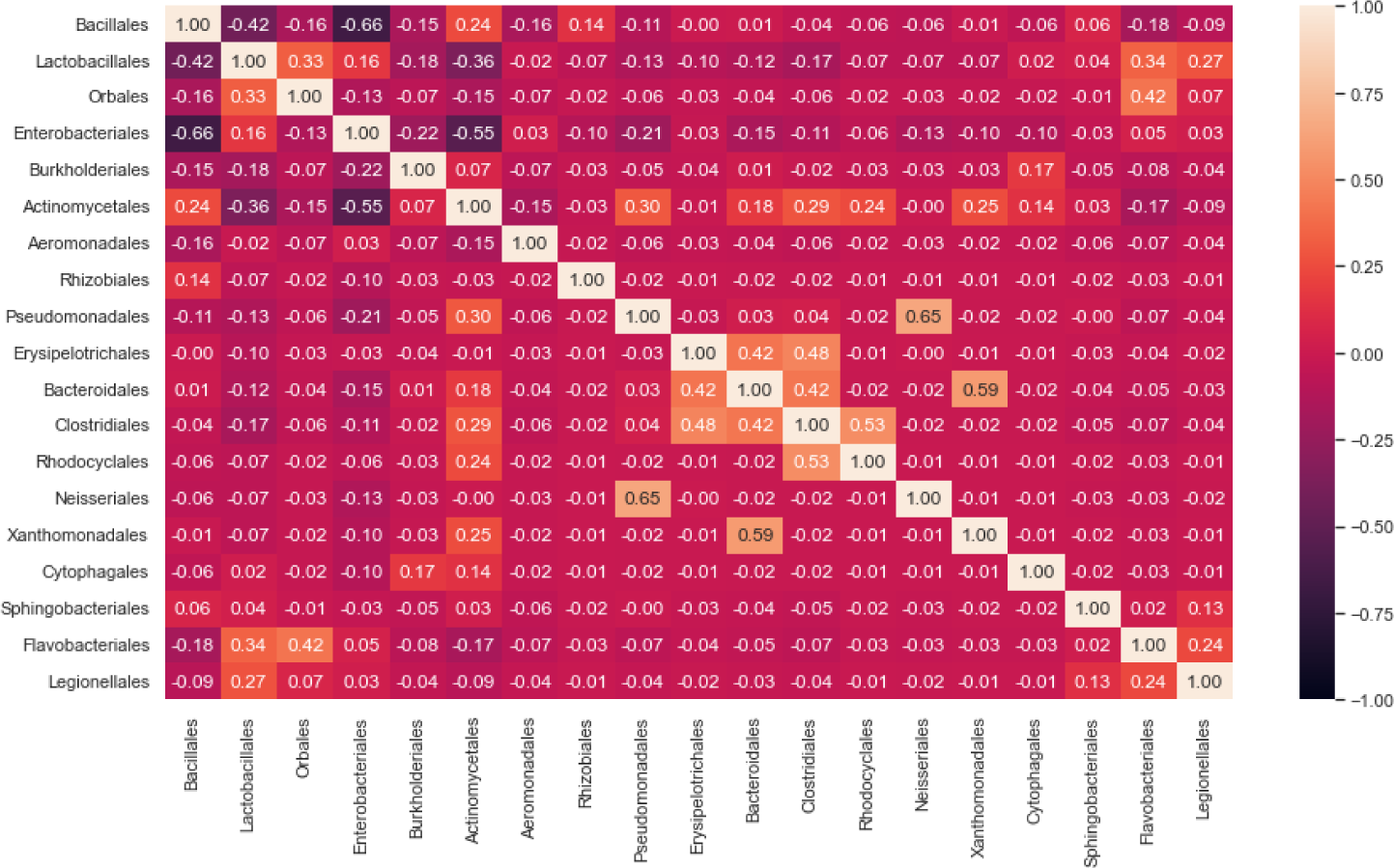
Correlation heatmap for all the 19 families’ samples used in the experiment, which reveals a strong correlation among *Bacillales*, *Lactobacillales*, *Enterobacteriales* and *Actinomycetales*.

Some notable, cooperative relationships exist amongst the data, particularly between *Orbales* and *Lactobacillales*, *Flavobacteriales* and *Lactobacillales*, and *Actinomycetales* and *Pseudomonadales*, having correlation coefficients of 0.33, 0.34 and 0.30, respectively. Interestingly, the correlation coefficient between *Lactobacillales* and *Enterobacteriales*, which were observed to be the dominant bacterial families present in the adult developmental stages, was only measured to be 0.16. Likewise, the correlation coefficient between *Actinomycetales* and *Bacillales*, which were observed to be dominant in the pupae developmental stage, was only measured to be 0.24.

Our result on relative abundance aligns with previous studies. During the 5^th^ generation developmental stage, bacterial families consistently clustered together Majumder et al. (2020, 2022). One such family, *Enterobacteriales*, was found to have increased abundance in all developmental stages Majumder et al. (2022). We also observed that the pupae stage exhibited the lowest relative abundance levels, corresponding with findings from Majumder et al. (2020).

### 3.2 Performance of Intrafamilial Successor Prediction Machine Learning Model

This model aimed to predict the relative abundance of each bacterial family in Generation 5 using the past four generations as predictors. The output of the developed model only predicted that seven bacterial families should be present in Generation 5 of each developmental stage. Obtained from the leave-one-out cross-validation, the mean RMSE between the actual and predicted relative abundances of each bacterial family in Generation 5 is presented in Table 1. The largest errors were observed in the *Enterobacteriales* and *Burkholderiales* families, with the model getting the relative abundance of these families wrong by roughly 40%, on average. While not large, significant error was also observed in the prediction of *Aeromonodales* relative abundances, which was roughly 23% incorrect on average. Far less error was observed in the *Bacillales*, *Lactobacillales*, *Actinomycetales* and *Pseudomonadales* families with an average error of approximately 1%, 5%, 5% and 0.02% respectively. *Pseudomonadales* are known to play a role in terpenoid biosynthesis in organic samples and fluorobenzoate degradation in conventional sites Bartuv et al. (2023); Devpura et al. (2017); McDonald et al. (2019). They are sometimes considered the primary symbiont group in many terrestrial isopod populations Bredon et al. (2019); Dittmer and Bouchon (2018).

**Table 1.** RMSE *±* Standard Error for Intrafamilial Successor Prediction Model of the Relative Abundances of Bacterial Families.

| Family | Least Angle Regression | Random Forest | Elastic-Net | Lasso |
| --- | --- | --- | --- | --- |
| *Bacillales* | 1.06 ± 0.31 | 0.67 ± 0.43 | 0.60 ± 0.44 | 0.58 ± 0.45 |
| *Lactobacillales* | 6.89 ± 1.84 | 5.05 ± 1.70 | 3.64 ± 1.44 | 3.80 ± 1.56 |
| *Enterobacteriales* | 45.59 ± 8.14 | 48.74 ± 9.25 | 38.89 ± 7.78 | 41.16 ± 8.77 |
| *Burkholderiales* | 38.14 ± 11.01 | 36.42 ± 11.82 | 46.64 ± 9.36 | 42.84 ± 15.75 |
| *Actinomycetales* | 6.10 ± 1.76 | 5.49 ± 2.17 | 4.36 ± 2.52 | 4.36 ± 2.52 |
| *Aeromonadales* | 22.85 ± 5.09 | 23.0 ± 5.01 | 22.85 ± 5.09 | 22.85 ± 5.09 |
| *Pseudomonadales* | 0.02 ± 0.01 | 0.02 ± 0.01 | 0.02 ± 0.01 | 0.02 ± 0.01 |
| Mean | 17.24 ± 4.02 | 17.05 ± 4.34 | 16.71 ± 3.81 | 16.52 ± 4.88 |

The mean RMSE was consistent across all methods, with the best-performing method, Lasso, making an average error of 16.52%, and the worst-performing method, Least Angle Regression, making an average error of 17.24%. Similarly, the Elastic-Net algorithm made an average error of 16.71%, while the Random Forest method made an average error of 17.05%.

The predicted relative abundances can be graphed against the actual values to better characterise this model’s performance and understand the source of these errors. A figure of predicted versus the actual relative abundance of the *Lactobacillales* family within the adult male developmental stage for each of the four ML methods has been shown in the supplementary material (Fig. 7). The best-performing method for this particular case was Lasso, with an RMSE of 8.19. Across all methods, while it appears that the model correctly predicts higher values when the actual values increase, a much lower magnitude in the predicted relative abundances is observed compared to the actual values. This is much more noticeable in the Least Angle Regression output, where the predicted relative abundance does not exceed 10% when the actual values can get as large as 25%.

A similar comparison is presented in a figure in the supplementary material (Fig. 8), showing the predicted vs actual relative abundance of the *Enterobacteriales* family within the adult male developmental stage for each of the four ML methods. Lasso was the best-performing method for this particular case, with an RMSE of 22.22. For all of the methods used, the model could not successfully attribute higher predicted relative abundances with higher actual relative abundances, resulting in no clear relationships between predicted and actual values. The Elastic-Net and Lasso methods output an almost identical prediction, while the Random Forest method outputs a much higher range of relative abundances.

### 3.3 Performance of Interfamilial Quantitative Prediction Machine Learning Model

This model aimed to predict the amount of a given bacterial family in each sample using the amount of all other bacteria present in the sample as predictors. Obtained from the leave-one-out cross-validation, the mean Coefficient of Variance (CV) between the actual and predicted relative abundances of each bacterial family in Generation 5 is presented in Table 2. Across all ML methods, the model consistently predicted bacterial families with the least error were *Lactobacillales* and *Enterobacteriales*.

**Table 2.** Coefficient of Variance *±* Standard Error for Interfamilial Quantitative Prediction Model of the Relative Abundances of Bacterial Families.

| Family | Least Angle Regression | Random Forest | Elastic-Net | Lasso |
| --- | --- | --- | --- | --- |
| *Bacillales* | $1.58 \pm 0.12$ | $0.54 \pm 0.11$ | $1.35 \pm 0.19$ | $33.11 \pm 32.09$ |
| *Lactobacillales* | $1.20 \pm 0.08$ | $0.35 \pm 0.05$ | $0.94 \pm 0.10$ | $1.14 \pm 0.31$ |
| *Orbales* | $1.79 \pm 0.32$ | $0.85 \pm 0.24$ | $2.06 \pm 0.64$ | $1.31 \pm 0.33$ |
| *Enterobacteriales* | $1.02 \pm 0.04$ | $0.21 \pm 0.04$ | $0.90 \pm 0.05$ | $0.82 \pm 0.10$ |
| *Burkholderiales* | $1.89 \pm 0.36$ | $0.31 \pm 0.11$ | $1.82 \pm 0.37$ | $1.54 \pm 0.38$ |
| *Actinomycetales* | $1.47 \pm 0.18$ | $0.59 \pm 0.15$ | $0.86 \pm 0.18$ | $2.20 \pm 1.45$ |
| *Aeromonadales* | $1.81 \pm 0.33$ | $1.45 \pm 0.27$ | $1.80 \pm 0.33$ | $1.37 \pm 0.35$ |
| *Rhizobiales* | $2.00 \pm 0.99$ | $2.22 \pm 1.11$ | $2.00 \pm 1.00$ | $1.92 \pm 1.00$ |
| *Pseudomonadales* | $1.74 \pm 0.49$ | $0.95 \pm 0.37$ | $1.80 \pm 0.53$ | $2.22 \pm 0.87$ |
| *Erysipelotrichales* | $1.96 \pm 0.66$ | $2.06 \pm 0.76$ | $1.95 \pm 0.71$ | $1.69 \pm 0.72$ |
| *Bacteroidales* | $1.93 \pm 0.53$ | $1.35 \pm 0.55$ | $1.63 \pm 0.57$ | $1.41 \pm 0.57$ |
| *Clostridiales* | $1.82 \pm 0.33$ | $1.31 \pm 0.34$ | $1.40 \pm 0.32$ | $1.07 \pm 0.35$ |
| *Rhodocyclales* | $2.00 \pm 0.99$ | $1.69 \pm 1.18$ | $2.03 \pm 1.06$ | $1.80 \pm 1.04$ |
| *Neisseriales* | $1.95 \pm 0.76$ | $1.82 \pm 0.86$ | $2.06 \pm 0.77$ | $1.30 \pm 0.78$ |
| *Xanthomonadales* | $1.82 \pm 0.80$ | $1.83 \pm 1.00$ | $1.87 \pm 0.90$ | $1.62 \pm 0.87$ |
| *Cytophagales* | $2.00 \pm 0.99$ | $1.73 \pm 1.05$ | $2.00 \pm 0.99$ | $1.03 \pm 1.00$ |
| *Sphingobacteriales* | $1.66 \pm 0.41$ | $1.27 \pm 0.48$ | $1.45 \pm 0.41$ | $1.41 \pm 0.40$ |
| *Flavobacteriales* | $1.74 \pm 0.29$ | $0.88 \pm 0.28$ | $1.23 \pm 0.28$ | $1.23 \pm 0.28$ |
| Unknown | $1.84 \pm 0.38$ | $1.49 \pm 0.4$ | $1.28 \pm 0.37$ | $1.26 \pm 0.37$ |
| *Legionellales* | $1.91 \pm 0.59$ | $2.09 \pm 0.69$ | $1.85 \pm 0.59$ | $1.76 \pm 0.59$ |
| Mean | $1.47 \pm 0.48$ | $1.25 \pm 0.50$ | $1.61 \pm 0.52$ | $3.06 \pm 2.19$ |

The presence of *Lactobacillales* is a common factor that drives divergence in dietary treatment groups Klammsteiner et al. (2021). This group has been shown to improve pesticide resistance and help control gastrointestinal pathogens Trinder et al. (2015); Daisley et al. (2017). Organic waste that has been inoculated with *Lactobacillales* tends to yield higher biomass output and a better nutritional spectrum in black soldier fly larvae compared to waste amended with artificial feed Somroo et al. (2019). *Lactobacillales* also play a role in vitamin and cofactor metabolism McMullen et al. (2021). On the other hand, the *Enterobacteriales* family has been identified as an important biomarker for schizophrenia Zhuang et al. (2020); Shen et al. (2018); Nguyen et al. (2019). This family is known to produce short-chain fatty acids Zhuang et al. (2020), which likely play a central role in the microbiota-host crosstalk that regulates brain function and behaviour Caspani and Swann (2019); Van De Wouw et al. (2018). The families that the model consistently predicted with the highest error were *Rhizobiales*, *Rhodocyclales*, *Erysipelotrichales* and *Neisseriales*.

The Random Forest method was by far the most accurate in its predictions, with a mean CV of 1.25. The Least Angle Regression and Elastic-Net methods performed similarly, with a mean CV of 1.76 and 1.61, respectively. Due to the large inaccuracies in predicting the *Bacillales* family, the Lasso method performed worst, with a mean CV of 3.06.

### 3.4 Performance of Interfamilial Qualitative Prediction Machine Learning Model

This model aimed to predict the relative abundance of each bacterial family within a sample using binary information on the presence or absence of all bacterial families as predictors. Furthermore, this model aimed to build in further information detailing the generation and developmental stage from which the sample originated. A comparative analysis of the overall performance between the model with information limited strictly to the presence or absence of bacterial families and successive models with more information introduced is presented in Fig. 6, which presents the mean RMSE for each model. From this comparison, it can be seen that the mean RMSE for the model limited to information on the presence and absence of bacterial families is 7.14, while the introduction of information detailing which generation the sample originates from results in a slight increase in the mean RMSE to 7.30. Adding information about the developmental stage of the sample to the model leads to the biggest improvement in performance, which reduces the mean RMSE to 5.27. Only a slight reduction in the RMSE to 5.12 is observed when the model is fed with all available information.

**Fig. 6.**
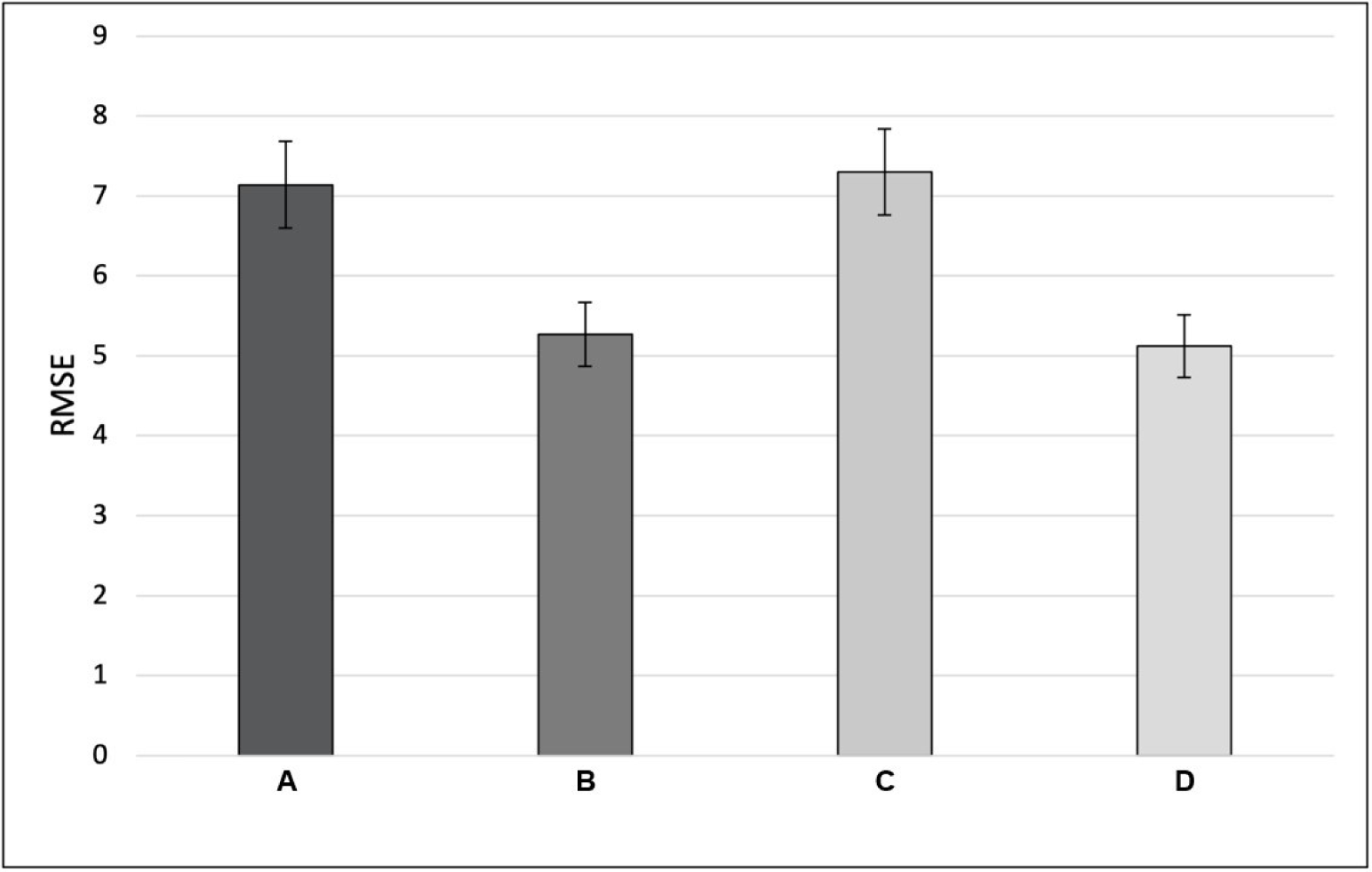
Mean RMSE between actual and predicted relative abundances for the different ML methods used in constructing the Interfamilial Qualitative Prediction model with error bars representing the standard error of the mean. Here, the first column “A” contains information about the presence of bacterial families, indicating whether a particular bacterial family is present in a given metagenomic sample. In the second column “B”, information regarding the developmental stage has been added along with the presence status. Similarly, in the third column “C”, additional generational information has been added along with information similar to the first column. Finally, the fourth column “D” contains all the information about a microbial family.

## 4 Discussion

### 4.1 Variance Between Different Generations of Laboratory-Reared Fruit Flies Leads to Significant Errors in Intrafamilial Models

The overarching aim of this research was to use metagenomic samples taken from fruit flies to analyse their microbiome composition and, consequently, understand whether different machine learning (ML) models could make predictions of the microbiome composition. Regarding the intrafamilial model developed in this research, the goal was to use this tool to observe how these microbiome compositions may change across different generations within a given developmental stage of flies. Using four different ML models, the relative abundances of seven different bacterial families were predicted for Generation 5 of each developmental stage, using the past four generations as predictors. By performing a comparative analysis of the mean RMSE between the actual and predicted relative abundances of the seven different bacterial families, no discernible difference was observed between the ML methods. This lack of diversity in performance amongst the methods is largely attributed to the poor performance of the overall model, with the relative abundance of some bacterial families being predicted in errors of close to 50% for all methods.

While similarly large errors are observed across several bacterial families, the reason for these errors can be attributed to a range of different sources. For example, the *Enterobacteriales* family had a mean RMSE between 39% and 48%; however, the average relative abundance of *Enterobacteriales* in Generation 5 amongst all developmental stages was approximately 60%. Furthermore, the range of *Enterobacteriales* relative abundance was the largest observed, with almost 100% relative abundance in Generation 5 of larvae but only 10% in Generation 5 of pupae. This high average and the large range of relative abundances lead to a significant error in prediction by the model, as no discernible pattern can be resolved amongst the highly variable outcomes. While this was the case for *Enterobacteriales*, the *Burkholderiales* family, which had a mean RMSE between 36% and 47%, was predicted in error due to its relative absence in all developmental stages, except the pupae stage.

Although the poor performance of this model is disappointing, the results reiterate previous observations made by Majumder et al. (2022), where they describe significant shifts in the overall microbiome composition between Generations 1 and 5. This large variance between generations inherently leads to considerable uncertainty in predictions made by the ML model, particularly noting that the changes between generations are completely different across developmental stages. Furthermore, for the data used in the development of this model, the maximum sample size within a generation was six, which limits the ability of the ML model to identify and build on any significant patterns that may exist within or between generations. As a result of the high variability in the microbiome composition between generations, coupled with the low sample size used to characterise these populations, a novel conclusion cannot be drawn from the results of this model, particularly regarding how the microbiome compositions may change across different generations. As described consistently throughout the literature, these microbiomes are highly complex systems that often display no discernible patterns in how they develop, change and evolve over time Australian Bureau of Agricultural Resource Economics and Sciences; Zhao et al. (2018). Moreover, within a given microbiome, there is a high amount of functional redundancy between bacterial families, whereby some bacteria can easily be replaced by others with similar functions without causing any changes to the fly’s physiology Moya and Ferrer (2016). As a result of this high complexity and plasticity within the gut microbiomes of fruit flies, models that can accurately predict the dynamics of these communities across different generations would require far more detail in the development of the ML algorithms used, alongside a much deeper pool of data to train these models with.

### 4.2 Random Forest Algorithm is More Appropriate in Characterising Interfamilial Models

Building on the observations made within the first model and towards achieving the overall goal of using metagenomic samples to analyse and predict the microbiome composition of fruit flies, the interfamilial model developed in this research was to characterise the relationships between different bacterial families. Across all ML methods, vastly different prediction patterns were observed, indicating that these relationships may be harder to decipher than anticipated. A comparative analysis of the mean Coefficient of Variance (CV) between the actual and predicted amounts of each bacterial family within all samples demonstrated that the Random Forest ML method was the most capable of producing predictions resembling the actual measurements. When the output of each method was compared to the actual number of bacterial families present, it was clear that Least Angle Regression, Elastic-Net and Lasso methods produced inaccurate predictions. Specifically, there was no resolvable relationship between the predictions generated by these methods and the actual observations in the samples. The Least Angle Regression model predictions aligned almost perfectly linearly with the actual values; however, the relationship was inverted. Furthermore, the range of values in which the Least Angle Regression model predicted the amount of bacteria should be present was far smaller than what was actually measured. The Elastic-Net and Lasso methods could output predictions that somewhat more accurately resembled the actual amounts of the bacterial families present within the samples; however, the performance of these methods remained very poor. While these three ML methods yielded no useful predictions, the Random Forest method produced some surprising results. This method yielded a somewhat proportional, linear relationship between predicted and actual amounts for most bacterial families. This is highlighted by Random Forest having the lowest mean CV, significantly outperforming the other three models. For some rarer bacterial families, this method tended to make more errors in prediction; however, more importantly, for almost all of the bacterial families, this method successfully predicted the correct range of the amounts of bacteria. While the model’s results signify that Random Forest methods are best suited to microbiome evaluation, the model’s accuracy still significantly limits its usefulness as a reliable predictive model in its current form. To accurately pinpoint the source of these errors, some rarer bacterial families could be removed from the dataset to allow the model to hone in on the more prevalent families.

Despite the positive results of this model, its usefulness in a wider context should be considered. The fundamental basis of this model implies that there is some existing information on the composition of a gut microbiome that can be used to predict an otherwise unknown amount of a selected bacterial family. To influence the physiology and behaviour of fruit flies and their impact on important crops, it may be possible to introduce certain bacterial families to create significant shifts in microbiome composition. As a result, an ML model such as that generated in this study may be able to be fed information on the existing composition of a gut microbiome and provide indications as to how bacterial family candidates may propagate amongst the system. Obviously, this would require the model to be trained on systems where the candidate bacterial family is already present to obtain any existing relationships.

### 4.3 Predictions of the Overall Microbiome Composition Are Largely Driven by the Developmental Stage

The final model within this study aimed to understand whether the complete microbiome composition of fruit flies could be predicted, only utilising information that detailed which bacterial families were present and which generation and developmental stage the samples originated from. A clear comparison between how such information informs the model was drawn by systematically integrating these elements into the predictive model. The most basic of these models, in which the presence or absence of bacteria was the only information fed in, was able to predict the overall composition of samples with a relatively low amount of error. The performance of this basic model was largely limited to predicting the correct types of bacteria present rather than the correct relative abundances. Upon the introduction of information detailing which generation the samples originated from, no discernible difference in the model’s performance was observed. These results can be interpreted in two ways: (*i*) the generation to which a fruit fly belongs does not significantly contribute to its overall microbiome composition; or (*ii*) the variance amongst the gut microbiomes within and between each generation is so large that no reasonable pattern can be drawn.

The introduction of indicators for which developmental stage the samples belonged yielded the largest increase in performance of the model, with a reduction to the lowest levels of prediction error observed throughout any of the models developed in this study. This increase in performance from the base model was largely attributed to the readjustment of the correctly predicted bacterial family relative abundances towards the true values. Furthermore, introducing this information often resulted in the model correctly removing bacterial families that the base model predicted should be present but were, in fact, absent from the actual samples.

Despite these promising results, there were instances where the iterations of the model were unable to accurately predict the overall composition of some microbe samples. They often overestimated the amounts of particular bacterial families and included bacterial families that were entirely absent from the actual measurements. While the source of these errors is hidden within the cryptic patterns between the relative abundances of different bacterial families, the results further signify and reiterate the highly complex nature of these systems.

To enhance the proposed method further, one promising approach could be transfer learning, yielding more positive results in classifying microbial communities Chong et al. (2022). Moreover, more curated data normalisation and other feature engineering techniques should be applied to maintain a balance between the range of the data. This would help reduce the weight biases of the ML model. Additionally, establishing a uniform dataset based on lab-grown species as a standard would facilitate better comparison of the models’ performance. Lastly, introducing Explainable AI (XAI) can make the models much more readable, ultimately aiding in translating our findings for other gut microbiome studies.

### 4.4 Comparison with Existing Literature

In the context of animal gut microbiome comparison, the results from Turner et al. (2022) were surprising, with an accuracy rate of over 90% (equivalent to an error percentage of 10%). Experiments conducted by Pietrucci et al. (2020) on the microbiota dysbiosis in Parkinson’s disease yielded an AUC score of 0.8 and an accuracy score of 71% using the Random Forest algorithm. Another study conducted by Su et al. (2022) in Hong Kong produced an accuracy of 83.3% and AUC scores ranging from 0.90 to 0.99. Although it is challenging to compare all of these studies on a common ground due to variations in their methodologies, on average, we can say that each of these models performs with an error rate somewhere between 10% and 30%. This indicates a high variance in the models’ prediction capability compared to our proposed solution. A summary of the outcomes from existing literature compared to ours is shown in Table 3.

**Table 3.**
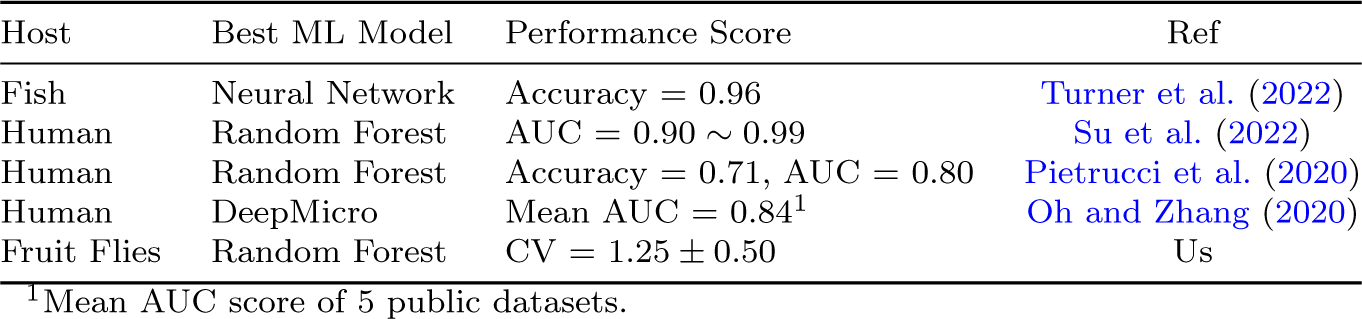
ML Model Performance Comparison With Existing Literature.

## 5 Conclusion

This study aimed to use metagenomic samples taken from fruit flies (tephritidae) to characterise their microbiome composition. Furthermore, this study aimed to use this knowledge to observe how these microbiome compositions change with the development of flies across their lifespan and between generations. The second key aim of this research was to understand whether the complete microbiome composition of fruit flies or elements within this microbiome could be predicted using simple machine learning (ML) models. Extending on this, a further aim was to understand which ML models best predict systems such as gut microbiomes. The first of these models did not perform well and could not accurately predict the relative abundances of bacterial families in a final generation. This was partly due to the large amount of variance between the gut microbiome composition of different generations of flies within the data used. The second model’s performance was promising and demonstrated that simple ML models can be built around the Random Forest method to predict gut microbiomes with reasonable accuracy. The final model was the most successful, able to predict the overall composition of gut microbiomes with significant accuracy. The overall results of this study firstly demonstrate how complex these dynamic systems are but also signify that more computationally efficient methods can be developed to determine microbiome compositions in a more practical way. The absence of simple ML models performing these types of analyses within the literature demonstrates the importance of the results within this study. It indicates that further parameter tuning within these models could yield greater accuracy. Although our model explored the fruit fly data as an example with our three proposed ML models, the methods are easily transferable to any other fruit fly or microbiome study.

## Data Availability

The datasets used during and/or analysed during the current study are publicly available in the NCBI database under Bioproject PRJNA717989 and are available at the following URL: https://www.ncbi.nlm.nih.gov/bioproject/?term=PRJNA717989.

## Supplementary Materials

**Table 4.** Number of Samples Per Each Developmental Stage and Generation.

| Developmental Stage | Generation | No. of Samples | Developmental Stage | Generation | No. of Samples |
| --- | --- | --- | --- | --- | --- |
| Larvae | 1 | 3 | Adult Female | 1 | 6 |
|  | 2 | 6 |  | 2 | 5 |
|  | 3 | 6 |  | 3 | 6 |
|  | 4 | 6 |  | 4 | 5 |
|  | 5 | 6 |  | 5 | 6 |
| Pupae | 1 | 6 | Adult Male | 1 | 6 |
|  | 2 | 6 |  | 2 | 6 |
|  | 3 | 6 |  | 3 | 6 |
|  | 4 | 6 |  | 4 | 6 |
|  | 5 | 6 |  | 5 | 6 |

**Fig. 7.**
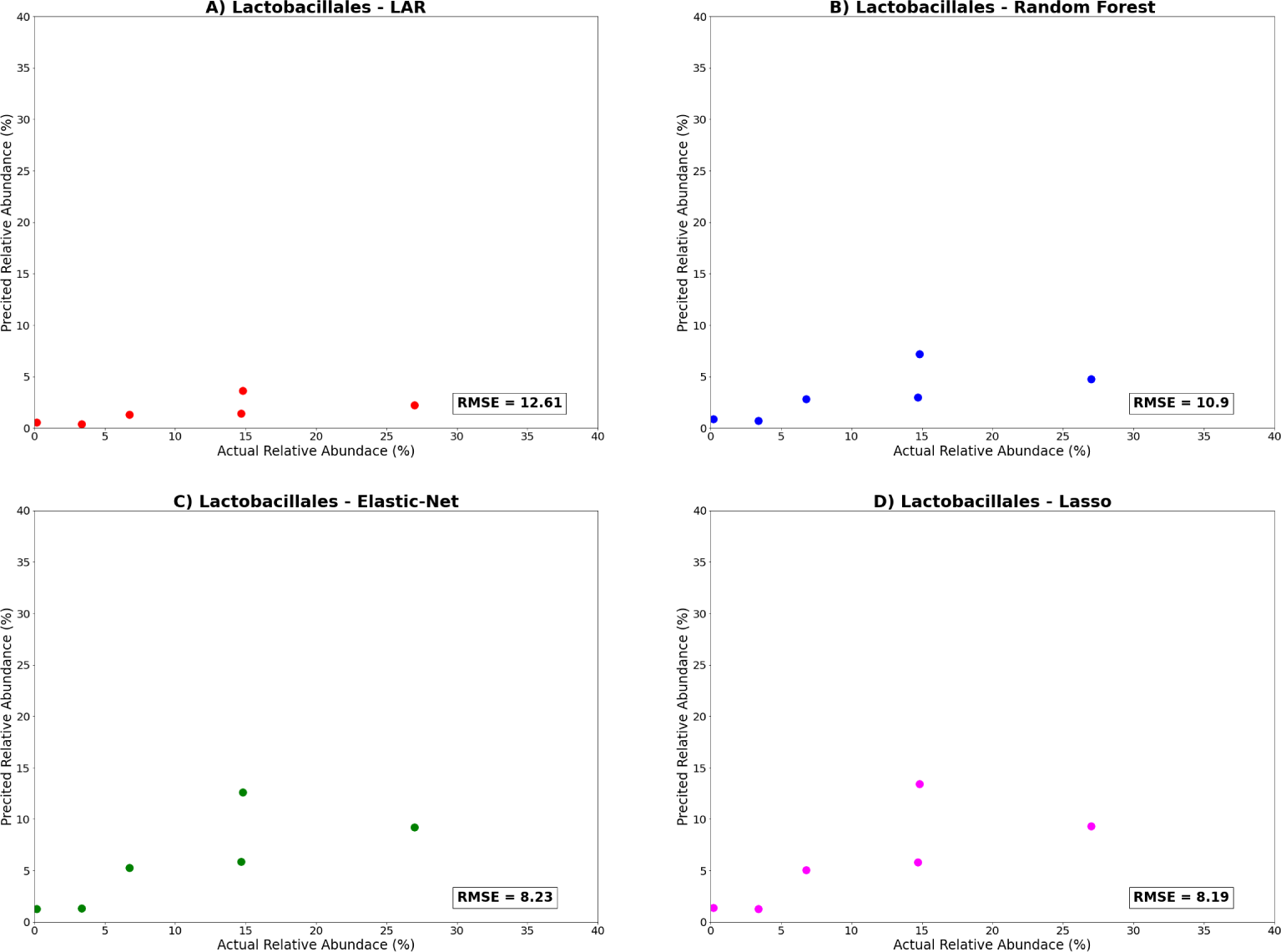
Intrafamilial model ML outputs predicting Generation 5 adult male *Lactobacillales* Relative Abundance for **(A)** Least Angle Regression, **(B)** Random Forest, **(C)** Elastic-Net and **(D)** Lasso.

**Fig. 8.**
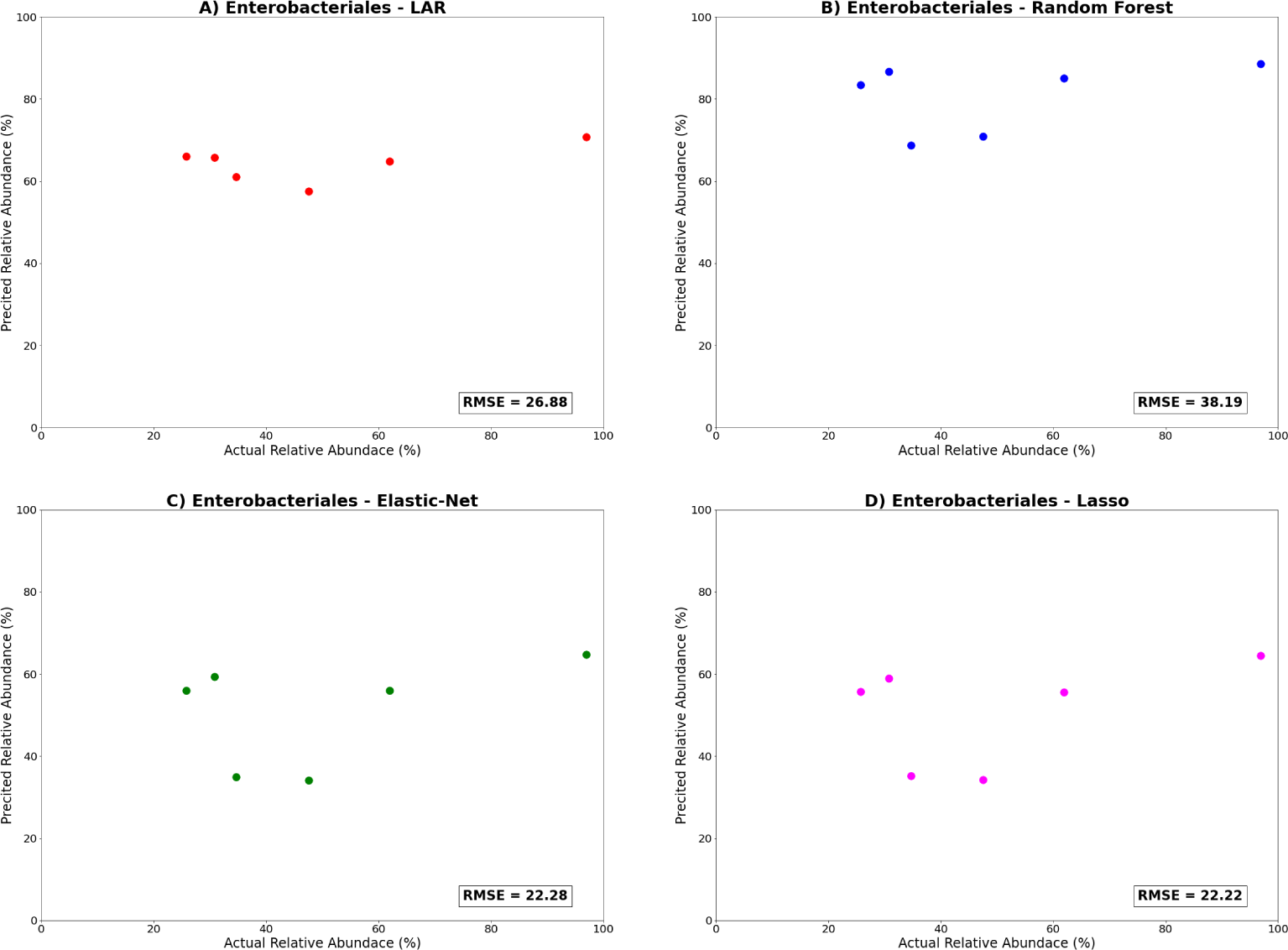
Intrafamilial model ML outputs predicting Generation 5 adult male *Enterobacteriales* Relative Abundance for **(A)** Least Angle Regression, **(B)** Random Forest, **(C)** Elastic-Net and **(D)** Lasso.

## References

Aidley, D.J.: Alternatives to insecticides. Science Progress (1933- ) 63(250), 293–303 (1976). Accessed 2023-01-17

Aktar, W., Sengupta, D., Chowdhury, A.: Impact of pesticides use in agriculture: their benefits and hazards. Interdisciplinary Toxicology 2(1), 1–12 (2009) 10.2478/v10102-009-0001-7

Australian Bureau of Agricultural Resource Economics and Sciences: Snapshot of Australian Agriculture 2022. https://www.agriculture.gov.au/abares/products/insights/snapshot-of-australian-agriculture-2022

Ami, E.B., Yuval, B., Jurkevitch, E.: Manipulation of the microbiota of mass-reared Mediterranean fruit flies Ceratitis capitata (Diptera: Tephritidae) improves sterile male sexual performance. The ISME Journal 4(1), 28–37 (2010) 10.1038/ismej.2009.82

Bakula, M.: The persistence of a microbial flora during postembryogenesis of Drosophila melanogaster. Journal of Invertebrate Pathology 14(3), 365–374 (1969) 10.1016/0022-2011(69)90163-3

Broderick, N.A., Buchon, N., Lemaitre, B.: Microbiota-Induced Changes in Drosophila melanogaster Host Gene Expression and Gut Morphology. mBio 5(3), 01117–14 (2014) 10.1128/mBio.01117-14

Bartuv, R., Berihu, M., Medina, S., Salim, S., Feygenberg, O., Faigenboim-Doron, A., Zhimo, V.Y., Abdelfattah, A., Piombo, E., Wisniewski, M., Freilich, S., Droby, S.: Functional analysis of the apple fruit microbiome based on shotgun metagenomic sequencing of conventional and organic orchard samples. Environmental Microbiology, 1462–292016353 (2023) 10.1111/1462-2920.16353

Bredon, M., Herran, B., Lheraud, B., Bertaux, J., Grève, P., Moumen, B., Bouchon, D.: Lignocellulose degradation in isopods: new insights into the adaptation to terrestrial life. BMC Genomics 20(1), 462 (2019) 10.1186/s12864-019-5825-8

Brown, C.E.: Coefficient of Variation. In: Brown, C.E. (ed.) Applied Multivariate Statistics in Geohydrology and Related Sciences, pp. 155–157. Springer, Berlin, Heidelberg (1998). 10.1007/978-3-642-80328-4 13

Caltagirone, L.E., Doutt, R.L.: The History of the Vedalia Beetle Importation to California and its Impact on the Development of Biological Control. Annual Review of Entomology 34(1), 1–16 (1989) 10.1146/annurev.en.34.010189.000245

Caspani, G., Swann, J.: Small talk: microbial metabolites involved in the signaling from microbiota to brain. Current Opinion in Pharmacology 48, 99–106 (2019) 10.1016/j.coph.2019.08.001

Chong, H., Zha, Y., Yu, Q., Cheng, M., Xiong, G., Wang, N., Huang, X., Huang, S., Sun, C., Wu, S., Chen, W.-H., Coelho, L.P., Ning, K.: EXPERT: transfer learning-enabled context-aware microbial community classification. Briefings in Bioinformatics 23(6), 396 (2022) 10.1093/bib/bbac396

Dittmer, J., Bouchon, D.: Feminizing Wolbachia influence microbiota composition in the terrestrial isopod Armadillidium vulgare. Scientific Reports 8(1), 6998 (2018) 10.1038/s41598-018-25450-4

Devpura, N., Jain, K., Patel, A., Joshi, C.G., Madamwar, D.: Metabolic potential and taxonomic assessment of bacterial community of an environment to chronic industrial discharge. International Biodeterioration & Biodegradation 123, 216–227 (2017) 10.1016/j.ibiod.2017.06.011

Daisley, B.A., Trinder, M., McDowell, T.W., Welle, H., Dube, J.S., Ali, S.N., Leong, H.S., Sumarah, M.W., Reid, G.: Neonicotinoid-induced pathogen susceptibility is mitigated by Lactobacillus plantarum immune stimulation in a Drosophila melanogaster model. Scientific Reports 7(1), 2703 (2017) 10.1038/s41598-017-02806-w

Furlan, L., Pozzebon, A., Duso, C., Simon-Delso, N., Sánchez-Bayo, F., Marchand, P.A., Codato, F., Lexmond, M., Bonmatin, J.-M.: An update of the Worldwide Integrated Assessment (WIA) on systemic insecticides. Part 3: alternatives to systemic insecticides. Environmental Science and Pollution Research 28(10), 11798–11820 (2021) 10.1007/s11356-017-1052-5

Glasl, B., Bourne, D.G., Frade, P.R., Thomas, T., Schaffelke, B., Webster, N.S.: Microbial indicators of environmental perturbations in coral reef ecosystems. Microbiome 7(1), 94 (2019) 10.1186/s40168-019-0705-7

Gould, A.L., Zhang, V., Lamberti, L., Jones, E.W., Obadia, B., Korasidis, N., Gavryushkin, A., Carlson, J.M., Beerenwinkel, N., Ludington, W.B.: Microbiome interactions shape host fitness. Proceedings of the National Academy of Sciences 115(51) (2018) 10.1073/pnas.1809349115

Hooper, L.V., Gordon, J.I.: Commensal Host-Bacterial Relationships in the Gut. Science 292(5519), 1115–1118 (2001) 10.1126/science.1058709

Hooper, L.V., Midtvedt, T., Gordon, J.I.: How host-microbial interactions shape the nutrient environment of the mammalian intestine. Annual Review of Nutrition 22(1), 283–307 (2002) 10.1146/annurev.nutr.22.011602.092259

Hernández Medina, R., Kutuzova, S., Nielsen, K.N., Johansen, J., Hansen, L.H., Nielsen, M., Rasmussen, S.: Machine learning and deep learning applications in microbiome research. ISME Communications 2(1), 1–7 (2022) 10.1038/s43705-022-00182-9

Hill, M.P., Macfadyen, S., Nash, M.A.: Broad spectrum pesticide application alters natural enemy communities and may facilitate secondary pest outbreaks. PeerJ 5, 4179 (2017) 10.7717/peerj.4179

Inamine, H., Ellner, S.P., Newell, P.D., Luo, Y., Buchon, N., Douglas, A.E.: Spatiotemporally Heterogeneous Population Dynamics of Gut Bacteria Inferred from Fecal Time Series Data. mBio 9(1), 01453–17 (2018) 10.1128/mBio.01453-17

Igbedioh, S.O.: Effects of Agricultural Pesticides on Humans, Animals, and Higher Plants in Developing Countries. Archives of Environmental Health: An International Journal 46(4), 218–224 (1991) 10.1080/00039896.1991.9937452

Kyritsis, G.A., Augustinos, A.A., Cáceres, C., Bourtzis, K.: Medfly Gut Microbiota and Enhancement of the Sterile Insect Technique: Similarities and Differences of Klebsiella oxytoca and Enterobacter sp. AA26 Probiotics during the Larval and Adult Stages of the VIENNA 8D53+ Genetic Sexing Strain. Frontiers in Microbiology 8, 2064 (2017) 10.3389/fmicb.2017.02064

Kyritsis, G.A., Augustinos, A.A., Ntougias, S., Papadopoulos, N.T., Bourtzis, K., Cáceres, C.: Enterobacter sp. AA26 gut symbiont as a protein source for Mediterranean fruit fly mass-rearing and sterile insect technique applications. BMC Microbiology 19(S1), 288 (2019) 10.1186/s12866-019-1651-z

Kirsch, P.: Pheromones: Their potential role in control of agricultural insect pests. American Journal of Alternative Agriculture 3(2-3), 83–97 (1988) 10.1017/S0889189300002241

Kumar, M., Ji, B., Zengler, K., Nielsen, J.: Modelling approaches for studying the microbiome. Nature Microbiology 4(8), 1253–1267 (2019) 10.1038/s41564-019-0491-9

Khomich, M., Måge, I., Rud, I., Berget, I.: Analysing microbiome intervention design studies: Comparison of alternative multivariate statistical methods. PLOS ONE 16(11), 0259973 (2021) 10.1371/journal.pone.0259973

Klammsteiner, T., Walter, A., Bogataj, T., Heussler, C.D., Stres, B., Steiner, F.M., Schlick-Steiner, B.C., Insam, H.: Impact of Processed Food (Canteen and Oil Wastes) on the Development of Black Soldier Fly (Hermetia illucens) Larvae and Their Gut Microbiome Functions. Frontiers in Microbiology 12, 619112 (2021) 10.3389/fmicb.2021.619112

Little, A.S., Light, S.H.: Stickland metabolism in the gut. Nature Microbiology 7(5), 603–604 (2022) 10.1038/s41564-022-01115-x

McMullen, J.G., Bueno, E., Blow, F., Douglas, A.E.: Genome-Inferred Correspondence between Phylogeny and Metabolic Traits in the Wild *Drosophila* Gut Microbiome. Genome Biology and Evolution 13(8), 127 (2021) 10.1093/gbe/evab127

Moya, A., Ferrer, M.: Functional redundancy-induced stability of gut microbiota subjected to disturbance. Trends in Microbiology 24(5), 402–413 (2016) 10.1016/j.tim.2016.02.002. Accessed 2023-02-15

Matos, R.C., Leulier, F.: Lactobacilli-Host mutualism: “learning on the fly”. Microbial Cell Factories 13(Suppl 1), 6 (2014) 10.1186/1475-2859-13-S1-S6

Majumder, R., Sutcliffe, B., Adnan, S.M., Mainali, B., Dominiak, B.C., Taylor, P.W., Chapman, T.A.: Artificial Larval Diet Mediates the Microbiome of Queensland Fruit Fly. Frontiers in Microbiology 11, 576156 (2020) 10.3389/fmicb.2020.576156

Majumder, R., Taylor, P.W., Chapman, T.A.: Dynamics of the Queensland Fruit Fly Microbiome through the Transition from Nature to an Established Laboratory Colony. Microorganisms 10(2), 291 (2022) 10.3390/microorganisms10020291

Mandal, S., Van Treuren, W., White, R.A., Eggesbø, M., Knight, R., Peddada, S.D.: Analysis of composition of microbiomes: a novel method for studying microbial composition. Microbial Ecology in Health & Disease 26(0) (2015) 10.3402/mehd.v26.27663

Michel-Mata, S., Wang, X., Liu, Y., Angulo, M.T.: Predicting microbiome compositions from species assemblages through deep learning. iMeta 1(1) (2022) 10.1002/imt2.3

McDonald, R.C., Watts, J.E.M., Schreier, H.J.: Effect of Diet on the Enteric Microbiome of the Wood-Eating Catfish Panaque nigrolineatus. Frontiers in Microbiology 10, 2687 (2019) 10.3389/fmicb.2019.02687

Nguyen, T.T., Kosciolek, T., Maldonado, Y., Daly, R.E., Martin, A.S., McDonald, D., Knight, R., Jeste, D.V.: Differences in gut microbiome composition between persons with chronic schizophrenia and healthy comparison subjects. Schizophrenia Research 204, 23–29 (2019) 10.1016/j.schres.2018.09.014

Nile, A.S., Kwon, Y.D., Nile, S.H.: Horticultural oils: possible alternatives to chemical pesticides and insecticides. Environmental Science and Pollution Research 26(21), 21127–21139 (2019) 10.1007/s11356-019-05509-z

Oh, M., Zhang, L.: DeepMicro: deep representation learning for disease prediction based on microbiome data. Scientific Reports 10(1), 6026 (2020) 10.1038/s41598-020-63159-5

Palmer, C., Bik, E.M., DiGiulio, D.B., Relman, D.A., Brown, P.O.: Development of the Human Infant Intestinal Microbiota. PLoS Biology 5(7), 177 (2007) 10.1371/journal.pbio.0050177

Pavlidi, N., Kampouraki, A., Tseliou, V., Wybouw, N., Dermauw, W., Roditakis, E., Nauen, R., Van Leeuwen, T., Vontas, J.: Molecular characterization of pyrethroid resistance in the olive fruit fly Bactrocera oleae. Pesticide Biochemistry and Physiology 148, 1–7 (2018) 10.1016/j.pestbp.2018.03.011

Pareek, A., Meena, B., Sharma, S., Tetarwal, M., Kalyan, R., Meena, B.: Impact of climate change on insect pests and their management strategies. Climate change and sustainable agriculture, 253–286 (2017)

Pasolli, E., Truong, D.T., Malik, F., Waldron, L., Segata, N.: Machine Learning Meta-analysis of Large Metagenomic Datasets: Tools and Biological Insights. PLOS Computational Biology 12(7), 1004977 (2016) 10.1371/journal.pcbi.1004977

Pietrucci, D., Teofani, A., Unida, V., Cerroni, R., Biocca, S., Stefani, A., Desideri, A.: Can Gut Microbiota Be a Good Predictor for Parkinson’s Disease? A Machine Learning Approach. Brain Sciences 10(4), 242 (2020) 10.3390/brainsci10040242

Reddy, V., Devi, M.J., Anbumozhi, V.: Ensuring food and nutritional security in the face of disasters and climate change: what is the adaptive solution. Towards a Resilient ASEAN Volume 1: Disasters, Climate Change, and Food Security: Supporting ASEAN Resilience, 290–330 (2019)

Ridley, E.V., Wong, A.C.-N., Westmiller, S., Douglas, A.E.: Impact of the Resident Microbiota on the Nutritional Phenotype of Drosophila melanogaster. PLOS ONE 7(5), 36765 (2012) 10.1371/journal.pone.0036765

Sundström, J.F., Albihn, A., Boqvist, S., Ljungvall, K., Marstorp, H., Martiin, C., Nyberg, K., Vågsholm, I., Yuen, J., Magnusson, U.: Future threats to agricultural food production posed by environmental degradation, climate change, and animal and plant diseases – a risk analysis in three economic and climate settings. Food Security 6(2), 201–215 (2014) 10.1007/s12571-014-0331-y

Steinigeweg, C., Alkassab, A.T., Erler, S., Beims, H., Wirtz, I.P., Richter, D., Pistorius, J.: Impact of a Microbial Pest Control Product Containing Bacillus thuringiensis on Brood Development and Gut Microbiota of Apis mellifera Worker Honey Bees. Microbial Ecology (2022) 10.1007/s00248-022-02004-w

Sultana, S., Baumgartner, J.B., Dominiak, B.C., Royer, J.E., Beaumont, L.J.: Impacts of climate change on high priority fruit fly species in Australia. PLOS ONE 15(2), 0213820 (2020) 10.1371/journal.pone.0213820

Schulz, N., Belheouane, M., Dahmen, B., Ruan, V.A., Specht, H.E., Dempfle, A., Herpertz-Dahlmann, B., Baines, J.F., Seitz, J.: Gut microbiota alteration in adolescent anorexia nervosa does not normalize with short-term weight restoration. International Journal of Eating Disorders 54(6), 969–980 (2021) 10.1002/eat.23435

Schnepf, E., Crickmore, N., Van Rie, J., Lereclus, D., Baum, J., Feitelson, J., Zeigler, D.R., Dean, D.H.: *Bacillus thuringiensis* and Its Pesticidal Crystal Proteins. Microbiology and Molecular Biology Reviews 62(3), 775–806 (1998) 10.1128/MMBR.62.3.775-806.1998

Savary, S., Ficke, A., Aubertot, J.-N., Hollier, C.: Crop losses due to diseases and their implications for global food production losses and food security. Food Security 4(4), 519–537 (2012) 10.1007/s12571-012-0200-5

Sivakala, K.K., Jose, P.A., Matan, O., Zohar-Perez, C., Nussinovitch, A., Jurkevitch, E.: *In vivo* predation and modification of the Mediterranean fruit fly *Ceratitis capitata* (Wiedemann) gut microbiome by the bacterial predator *Bdellovibrio bacteriovorus*. Journal of Applied Microbiology 131(6), 2971–2980 (2021) 10.1111/jam.15170

Su, Q., Liu, Q., Lau, R.I., Zhang, J., Xu, Z., Yeoh, Y.K., Leung, T.W.H., Tang, W., Zhang, L., Liang, J.Q.Y., Yau, Y.K., Zheng, J., Liu, C., Zhang, M., Cheung, C.P., Ching, J.Y.L., Tun, H.M., Yu, J., Chan, F.K.L., Ng, S.C.: Faecal microbiome-based machine learning for multi-class disease diagnosis. Nature Communications 13(1), 6818 (2022) 10.1038/s41467-022-34405-3

Somroo, A.A., Ur Rehman, K., Zheng, L., Cai, M., Xiao, X., Hu, S., Mathys, A., Gold, M., Yu, Z., Zhang, J.: Influence of Lactobacillus buchneri on soybean curd residue co-conversion by black soldier fly larvae (Hermetia illucens) for food and feedstock production. Waste Management 86, 114–122 (2019) 10.1016/j.wasman.2019.01.022

Savary, S., Willocquet, L., Pethybridge, S.J., Esker, P., McRoberts, N., Nelson, A.: The global burden of pathogens and pests on major food crops. Nature Ecology & Evolution 3(3), 430–439 (2019) 10.1038/s41559-018-0793-y

Shen, Y., Xu, J., Li, Z., Huang, Y., Yuan, Y., Wang, J., Zhang, M., Hu, S., Liang, Y.: Analysis of gut microbiota diversity and auxiliary diagnosis as a biomarker in patients with schizophrenia: A cross-sectional study. Schizophrenia Research 197, 470–477 (2018) 10.1016/j.schres.2018.01.002

Trinder, M., Bisanz, J.E., Burton, J.P., Reid, G.: Probiotic lactobacilli: a potential prophylactic treatment for reducing pesticide absorption in humans and wildlife. Beneficial Microbes 6(6), 841–847 (2015) 10.3920/BM2015.0022

Turner, J.W., Cheng, X., Saferin, N., Yeo, J.-Y., Yang, T., Joe, B.: Gut microbiota of wild fish as reporters of compromised aquatic environments sleuthed through machine learning. Physiological Genomics 54(5), 177–185 (2022) 10.1152/physiolgenomics.00002.2022

Tang, W., Wilkening, J., Desai, N., Gerlach, W., Wilke, A., Meyer, F.: A scalable data analysis platform for metagenomics. In: 2013 IEEE International Conference on Big Data, pp. 21–26 (2013). 10.1109/BigData.2013.6691723

Van De Wouw, M., Boehme, M., Lyte, J.M., Wiley, N., Strain, C., O’Sullivan, O., Clarke, G., Stanton, C., Dinan, T.G., Cryan, J.F.: Short-chain fatty acids: microbial metabolites that alleviate stress-induced brain-gut axis alterations: SCFAs alleviate stress-induced brain-gut axis alterations. The Journal of Physiology 596(20), 4923–4944 (2018) 10.1113/JP276431

Vontas, J., Hernández-Crespo, P., Margaritopoulos, J.T., Ortego, F., Feng, H.-T., Mathiopoulos, K.D., Hsu, J.-C.: Insecticide resistance in Tephritid flies. Pesticide Biochemistry and Physiology 100(3), 199–205 (2011) 10.1016/j.pestbp.2011.04.004

Wang, Z., Bovik, A.C.: Mean squared error: Love it or leave it? A new look at Signal Fidelity Measures. IEEE Signal Processing Magazine 26(1), 98–117 (2009) 10.1109/MSP.2008.930649

Yoon, B.W., Lim, S.-H., Shin, J.H., Lee, J.-W., Lee, Y., Seo, J.H.: Analysis of oral microbiome in glaucoma patients using machine learning prediction models. Journal of Oral Microbiology 13(1), 1962125 (2021) 10.1080/20002297.2021.1962125

Yang, F., Tomberlin, J.K., Jordan, H.R.: Starvation Alters Gut Microbiome in Black Soldier Fly (Diptera: Stratiomyidae) Larvae. Frontiers in Microbiology 12, 601253 (2021) 10.3389/fmicb.2021.601253

Yu, C., Zhou, Z., Liu, B., Yao, D., Huang, Y., Wang, P., Li, Y.: Investigation of trends in gut microbiome associated with colorectal cancer using machine learning. Frontiers in Oncology 13, 1077922 (2023) 10.3389/fonc.2023.1077922

Zhang, W., Jiang, F., Ou, J.: Global pesticide consumption and pollution: with china as a focus. Proceedings of the international academy of ecology and environmental sciences 1(2), 125 (2011)

Zhuang, Z., Yang, R., Wang, W., Qi, L., Huang, T.: Associations between gut microbiota and Alzheimer’s disease, major depressive disorder, and schizophrenia. Journal of Neuroinflammation 17(1), 288 (2020) 10.1186/s12974-020-01961-8

Zhao, X., Zhang, X., Chen, Z., Wang, Z., Lu, Y., Cheng, D.: The divergence in bacterial components associated with bactrocera dorsalis across developmental stages. Frontiers in Microbiology 9, 114 (2018) 10.3389/fmicb.2018.00114

